# The E3 ubiquitin ligase Pib1 regulates effective gluconeogenic shutdown in *S. cerevisiae*

**DOI:** 10.1101/673657

**Authors:** Vineeth Vengayil, Sunil Laxman

## Abstract

Cells use multiple mechanisms to regulate their metabolic states depending on changes in their nutrient environment. A well-known example is the response of cells to glucose availability. In *S. cerevisiae* cells growing in glucose-limited medium, the re-availability of glucose leads to the downregulation of gluconeogenesis, the activation of glycolysis, and robust ‘glucose repression’. However, our knowledge of the initial mechanisms mediating this glucose-dependent downregulation of the gluconeogenic transcription factors is incomplete. We used the gluconeogenic transcription factor Rds2 as a candidate with which to discover regulators of early events leading to glucose repression. Here, we identify a novel role for the E3 ubiquitin ligase Pib1 in regulating the stability and degradation of Rds2. Glucose addition to glucose-limited cells results in rapid ubiquitination of Rds2, followed by its proteasomal degradation. Through *in vivo* and *in vitro* experiments, we establish Pib1 as a ubiquitin E3 ligase that regulates Rds2 ubiquitination and stability. Notably, this Pib1 mediated Rds2 ubiquitination, followed by proteasomal degradation, is specific to the presence of glucose. Pib1 is required for complete glucose repression, and enables cells to optimally grow in competitive environments when glucose becomes re-available. Our results reveal the existence of a Pib1 E3-ubiquitin ligase mediated regulatory program that mediates glucose-repression when glucose availability is restored.

## Introduction

The uptake and utilization of nutrients are essential for cell growth and survival. Multiple metabolic pathways within cells make and break nutrients, and these pathways are tightly regulated depending on nutrient availability (1, 2). Since the nutrient environment of a cell is not constant, particularly for free-living microbes, cells rewire their metabolic pathways depending on the changes in nutrient availability (3–5). This ability of cells to rapidly and efficiently switch between different metabolic states is crucial for their survival and is closely coupled to the nutrient sensing machinery (6–8). Therefore, cells integrate multiple strategies to carry out efficient metabolic switching. Uderstanding these strategies are major areas of study.

*Saccharomyces cerevisiae* is an excellent model to understand conserved, general principles of metabolic state switching, due to the ease of controlling its metabolism (by altering nutrients provided), coupled with genetic and biochemical approaches to dissect regulatory mechanisms. A feature of *S. cerevisiae* metabolism is a preference for glucose as a carbon source, and where cells use fermentative metabolism (De Deken, 1966). This is the famous ‘Crabtree effect’, analogous to the Warburg effect in cancer cells, where cells preferentially use glucose over other available carbon sources, and minimize respiratory metabolism when glucose is present (De Deken, 1966; Diaz-Ruiz et al., 2011; Postma et al., 1989). Glucose availability, therefore, regulates a variety of cellular responses in yeast (12–14). Following glucose limitation, cells switch to a gluconeogenic state where alternate carbon sources are utilized, and upon glucose re-entry, cells switch back to a glycolytic state where alternate carbon source utilization is repressed (15, 16). Therefore, efficient glucose-induced catabolite repression is critical to ensure that upon glucose re-entry, gluconeogenesis is shut down (17–19). The initial responses occur immediately after glucose addition, through rapid changes in intracellular metabolite pools, driven by allosteric regulations and metabolic flux rewiring (20–22). Subsequently, the regulation of proteins that enforce the metabolic state switch, takes place by the interplay of transcriptional, translational, post-transcriptional and post-translational responses (23). Post-translation regulation (by signaling mediated events) allows rapid and dynamic regulation of protein levels and activity in response to a nutrient such as glucose (24). While we have a growing understanding of signaling events controlling cell growth with glucose as a carbon source (25–27), several gaps remain in our understanding of the ‘off-switches’ that enable effective glucose repression in cells.

In particular, we have a limited understanding of how regulated protein turnover controls metabolic state switching when cells encounter a new nutrient source. Our understanding of regulatory events in this condition is biased towards ‘classic’ signaling, through the activation of nutrient-responsive kinases and phosphatases (28–30). Alternate modes of regulation, including selective protein turnover in response to changing nutrients (as opposed to ‘starvation’) remain poorly studied. The ubiquitin-mediated proteasomal degradation is a major pathway of selective protein degradation in eukaryotes (31, 32), but the role of the ubiquitin-proteasomal system in regulating metabolic switching is poorly understood. Since target specificity of ubiquitination is achieved by E3 ubiquitin ligases, which bind and specifically target proteins for ubiquitin conjugation (33, 34), there must be distinct E3 ligases activated by unique nutrient cues, which then ubiquitinate their substrates. However, only a few studies identify roles of E3 ubiquitin ligases in the context of glucose-mediated metabolic switching. In yeast, upon glucose depletion, the E3 ligase Grr1 targets the phosphofructokinase (Pfk27) enzyme for degradation, thereby inhibiting glycolysis (35). In the context of catabolite (glucose) repression, the E3 ligase complex SCF_Ucc1_ regulates the degradation of the citrate synthase enzyme Cit2, thereby inhibiting the glyoxylate shunt (36). Further, the GID complex degrades the gluconeogenic enzymes Fbp1 and Pck1, in the presence of glucose (Santt et al., 2008; Schork et al., 2002). Apart from these, little is known about the role of E3 ligases in regulating effective gluconeogenic shutdown. Similar examples from mammalian cells are even rarer (39). Notably, these studies are limited to only the regulation of relevant metabolic enzymes following glucose addition. For a complete metabolic state switch, the transcription factors which regulate these enzyme transcripts must themselves be regulated. In yeast, Rds2, Cat8, and Sip4 are the transcription factors that regulate gluconeogenic enzyme transcripts during growth in glucose-limited conditions (1, 40–42). Although these transcription factors are well studied in cells growing in glucose limitation, how these factors are regulated after glucose becomes available has surprisingly not been addressed (Figure 1A).

**Figure 1:**
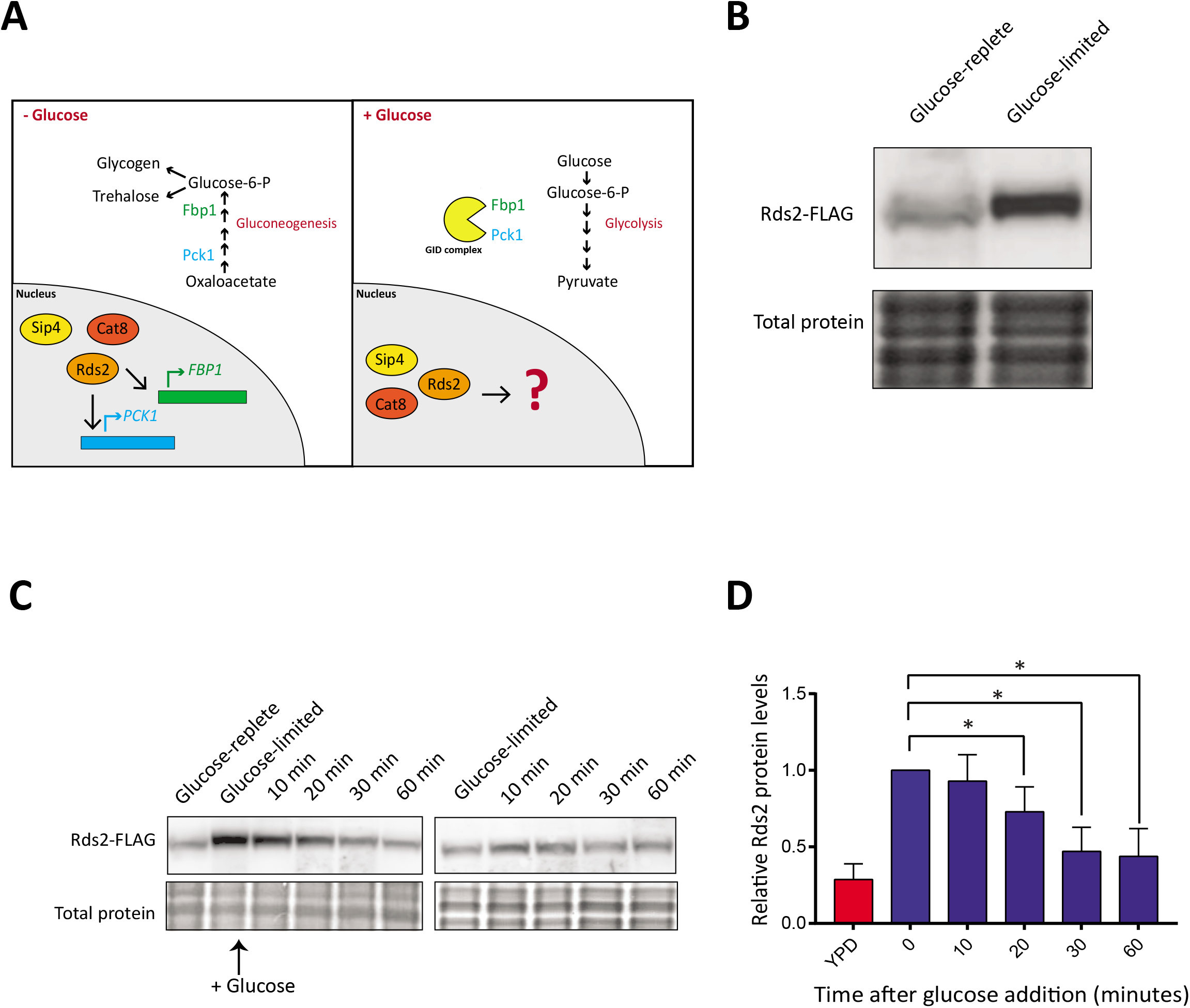
Glucose regulates Rds2 protein levels. A) An overview of the known transcriptional regulation of gluconeogenesis. In order to effectively switch to a glycolytic state after glucose re-entry, cells rapidly downregulate the gluconeogenic machinery. The gluconeogenic enzymes Pck1 and Fbp1 are regulated by a GID complex-mediated ubiquitination and proteasomal degradation. However, the regulation of the gluconeogenic transcription factors in the presence of glucose remains unknown. B) Glucose availability regulates amounts of the gluconeogenic activator Rds2. Cells were grown in glucose-replete or glucose-limited (glycerol/ethanol) medium, and Rds2 protein amounts were compared by Western blotting using anti-FLAG antibody (see Experimental Procedures). Coomassie-stained gel showing total protein was used as the loading control. A representative blot (from three independent experiments) is shown. C) Glucose addition results in a rapid reduction in Rds2 protein. Glucose was added to a final concentration of 3% to cells grown in glucose-limited medium. Cells were collected at different time points post glucose addition, and Rds2 amounts at each time were measured by Western blot using anti-FLAG antibody. Cells grown in glucose-limited medium (glycerol/ethanol) without the addition of glucose were used as a control. Loading control: Coomassie-stained gels showing total proteins. D) Quantification of the blot in (C). The relative Rds2 amounts were quantified with respect to the Rds2 amount in cells collected at 0 minutes in glucose-limited medium, using ImageJ. Data are displayed as means ± SD, n=3. *p<0.05, **p0.01, ***p<0.001.

In this study, we sought to identify novel regulators of effective glucose repression, by focusing on the gluconeogenic transcription factors when glucose becomes available. For this, we used the gluconeogenic transcription factor Rds2 as a candidate. Here, we discover that Rds2 is rapidly ubiquitinated and proteasomally degraded upon glucose addition. This is mediated by a specific E3 ubiquitin ligase – Pib1, in a glucose-dependent manner. Through *in vivo* and *in vitro* studies, we show a direct role of Pib1 in mediating glucose-dependent Rds2 ubiquitination and degradation. Finally, Pib1 is required for competitive cell growth upon glucose re-entry. Collectively, our study identifies a novel role for a previously obscure E3 ubiquitin ligase, in regulating gluconeogenic shutdown, and thereby facilitating effective glucose repression. More generally, our study exemplifies how E3-ubiquitin ligases can respond to changing nutrients, and serve as ‘on’ or ‘off’ switches to regulate cellular metabolic state.

## Results

### Glucose regulates Rds2 protein levels

The transcription factors Rds2, Cat8, and Sip4 regulate gluconeogenesis under conditions of glucose limitation, by activating the expression of key enzymes required for gluconeogenesis. However, the mechanisms by which these proteins are shut down following glucose re-entry are not well understood (Figure 1A). In order to understand the processes involved in glucose-mediated regulation of these transcription factors, we chose Rds2 as a candidate, to specifically understand regulation after glucose addition. Using yeast cells with Rds2 chromosomally tagged at the carboxy-terminus with a FLAG epitope, we monitored the steady-state levels of Rds2 in cells grown in high glucose medium (2% glucose) or low glucose medium (2% glycerol and ethanol) using Western blots. The amounts of Rds2 protein was substantially lower in cells in high glucose medium compared to low glucose medium (Figure 1B). This suggested an existing mechanism to regulate Rds2 protein levels depending on the availability of glucose. In order to understand the kinetics of Rds2 protein regulation when cells encounter glucose, we monitored Rds2 amounts over time in cells growing in glucose-limited medium (glycerol/ethanol as a carbon source) to which glucose was added. We observed a rapid decrease in Rds2 amounts after glucose addition (Figure 1C and Figure 1D), with a significant decrease observed as quickly as ~20 minutes after glucose addition (Figure 1D). After ~30 minutes of glucose addition, the amount of Rds2 protein was similar to that of the control (cells grown in 2% glucose medium). Further, no changes in Rds2 amounts were observed when glucose was not added (Figure 1C), reiterating that the change in Rds2 protein was glucose dependent. Overall, these results suggest that glucose addition activates a regulatory event that rapidly reduces the amount of Rds2 in cells.

### Glucose induces ubiquitination and proteasomal degradation of Rds2

Previous studies have identified roles of the protein degradation machinery in regulating the gluconeogenic enzymes Pck1 and Fbp1, in response to glucose addition (37). Protein degradation plays a role in rapid glucose-mediated catabolite inactivation of many proteins (Santt et al., 2008; Schüle et al., 2000). This is expected since decreases in protein amounts can be achieved swiftly via protein degradation, compared to changes in transcription or translation. Considering the decrease in Rds2 amounts within a few minutes after glucose re-entry, we hypothesized that glucose induces the regulated degradation of Rds2. Upon inspecting the amino acid sequence of Rds2, we found multiple lysine residues present in the protein (Figure 2A). Since ubiquitination (leading to protein degradation) occurs at the lysine residues of target proteins, we used multiple ubiquitination prediction programs (Wang 2017; Radivojac et al., 2011; Tung and Ho, 2008), to predict if these lysines were predicted to be ubiquitinated. We found that multiple lysine residues on Rds2 were strongly predicted to be likely ubiquitination sites (Figure 2A, Supplementary figure 1A). To test the possibility that Rds2 might indeed be ubiquitinated and degraded proteasomally, we first investigated if Rds2 was degraded by the proteasome. We treated cells with a proteasomal inhibitor-MG132, and monitored Rds2 protein amounts post glucose addition. Notably, Rds2 amounts remained constant after glucose addition (Figure 2B and Figure 2C), in contrast to our earlier observations in cells without MG132 (shown in Figure 1). This strongly suggested that glucose addition results in the proteasomal degradation of Rds2. We therefore asked whether Rds2 is conditionally ubiquitinated in response to glucose. For this, we added glucose to cells in the presence of MG132, immunopurified Rds2, and probed for poly-ubiquitin conjugates (by Western blotting). We observed ubiquitin conjugates only in Rds2 immunoprecipitates from glucose treated cells, but not in the control cells where glucose was not added (Figure 2D). This suggests that Rds2 is specifically ubiquitinated rapidly in cells, in a glucose dependent manner. Collectively these data indicate that Rds2 is ubiquitinated and proteasomally degraded when glucose becomes available in the medium.

**Figure 2:**
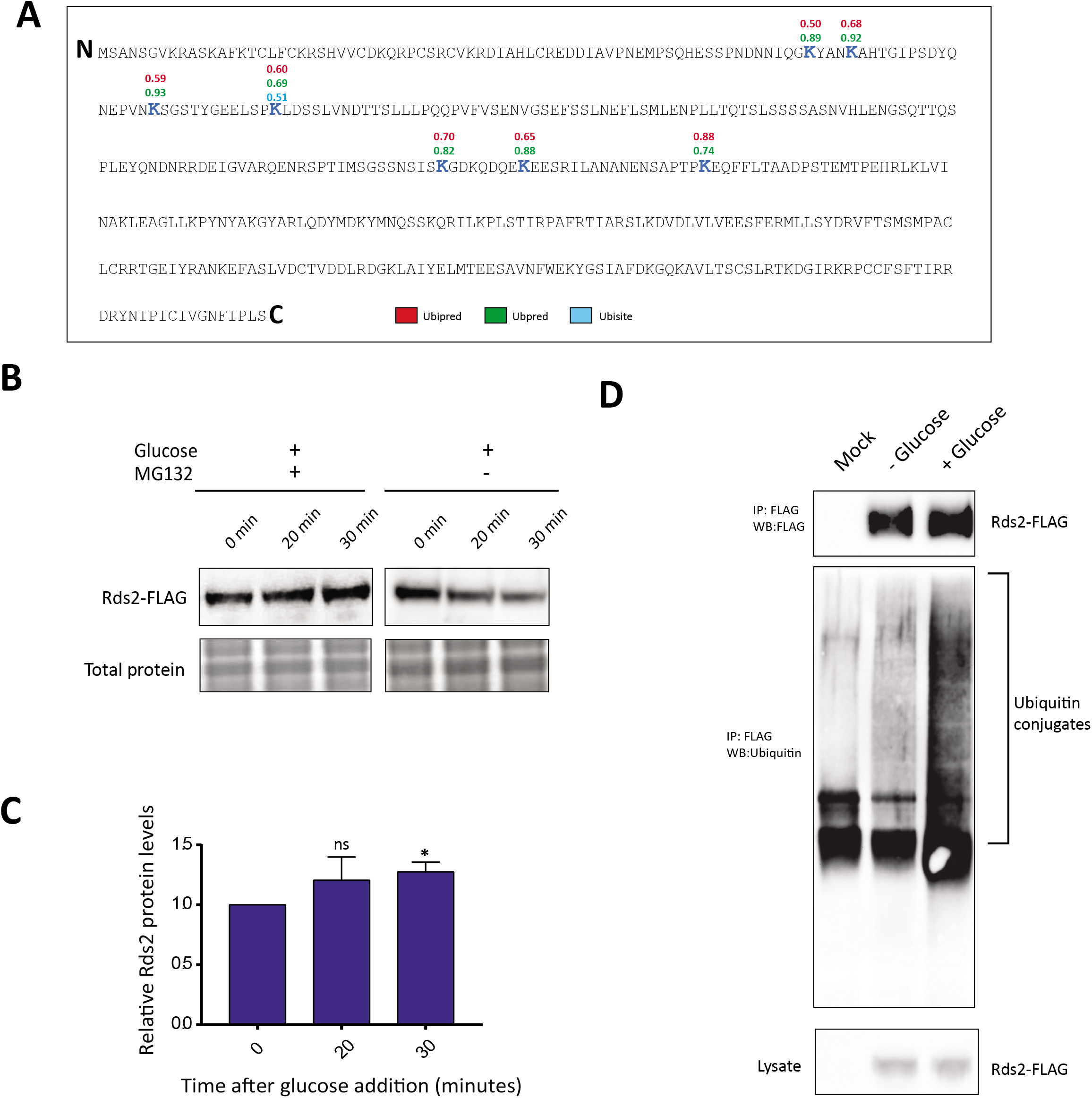
Glucose induces ubiquitination and proteasomal degradation of Rds2. A) Putative ubiquitination sites in Rs2p. We used three different ubiquitination prediction tools for identifying putative ubiquitin binding lysine residues in Rds2. The high score lysine residues (scores greater than 0.5) recognized by at least two of the three analysis software are shown in the schematic. Individual scores of all the lysine residues from the predictions are shown in Supplementary figure 1. B) Glucose-dependent degradation of Rds2 is mediated by the proteasome. Cells expressing Rds2-FLAG grown in glucose-limited medium were treated with glucose and MG132, and Rds2 amounts were measured by Western blot. Note: these cells also lack the multi-drug transporter Pdr5, to allow the MG132 treatment. Cells grown in glucose-limited medium with the addition of glucose without MG132 addition were used as the control. Coomassie-stained gels were used as loading control. C) Quantification of the blot in (B) using ImageJ. Data are displayed as means ± SD, n=3. *p<0.05, **p0.01, ***p<0.001. D) Glucose addition induces ubiquitination of Rds2. Rds2-FLAG was immunoprecipitated from glucose-treated and non-treated cells using anti-FLAG antibody. Rds2-ubiquitin conjugates were detected by western blotting using anti-ubiquitin antibody. A representative blot from at least three independent experiments is shown.

### The E3 ubiquitin ligase Pib1 interacts with and ubiquitinates Rds2

Ubiquitination is a multi-step process mediated by the coordinated action of three enzymes-E1 (ubiquitin activating), E2 (ubiquitin conjugating) and E3 (ubiquitin ligase). Ubiquitination substrate specificity depends on the E3 ubiquitin ligase, which selectively binds to and ubiquitinates the target protein (33). Therefore we hypothesized that in the presence of glucose, an unidentified E3 ubiquitin ligase should specifically ubiquitinate Rds2 and regulate its degradation (Figure 3A). We used YeastMine (47) to identify known interacting proteins of Rds2. This revealed, Pib1, a protein with E3 ligase activity, as an interacting protein partner of Rds2 (as seen in other high-throughput studies (48)). This putative interaction and possible regulatory function of Pib1 and Rds2 have remained unexplored. Therefore, to directly probe this putative interaction, we performed co-immunoprecipitation studies by immunopurified Pib1 (endogenously tagged at the carboxy-terminus with an HA epitope tag) and tested whether Rds2 associated with immunopurified Pib1. Notably, we performed these experiments using cells grown in (i) glucose-limited medium (2% glycerol and ethanol as a carbon source), (ii) without the addition of glucose, or (iii) 20 minutes after glucose addition. Strikingly, we observed a strong co-immunopurification of Rds2 with Pib1, only when Pib1 was isolated from cells minutes after glucose addition to the medium (Figure 3B). This strongly suggested that Pib1 interacts with Rds2, and specifically when glucose became available in the medium (Figure 3B).

**Figure 3:**
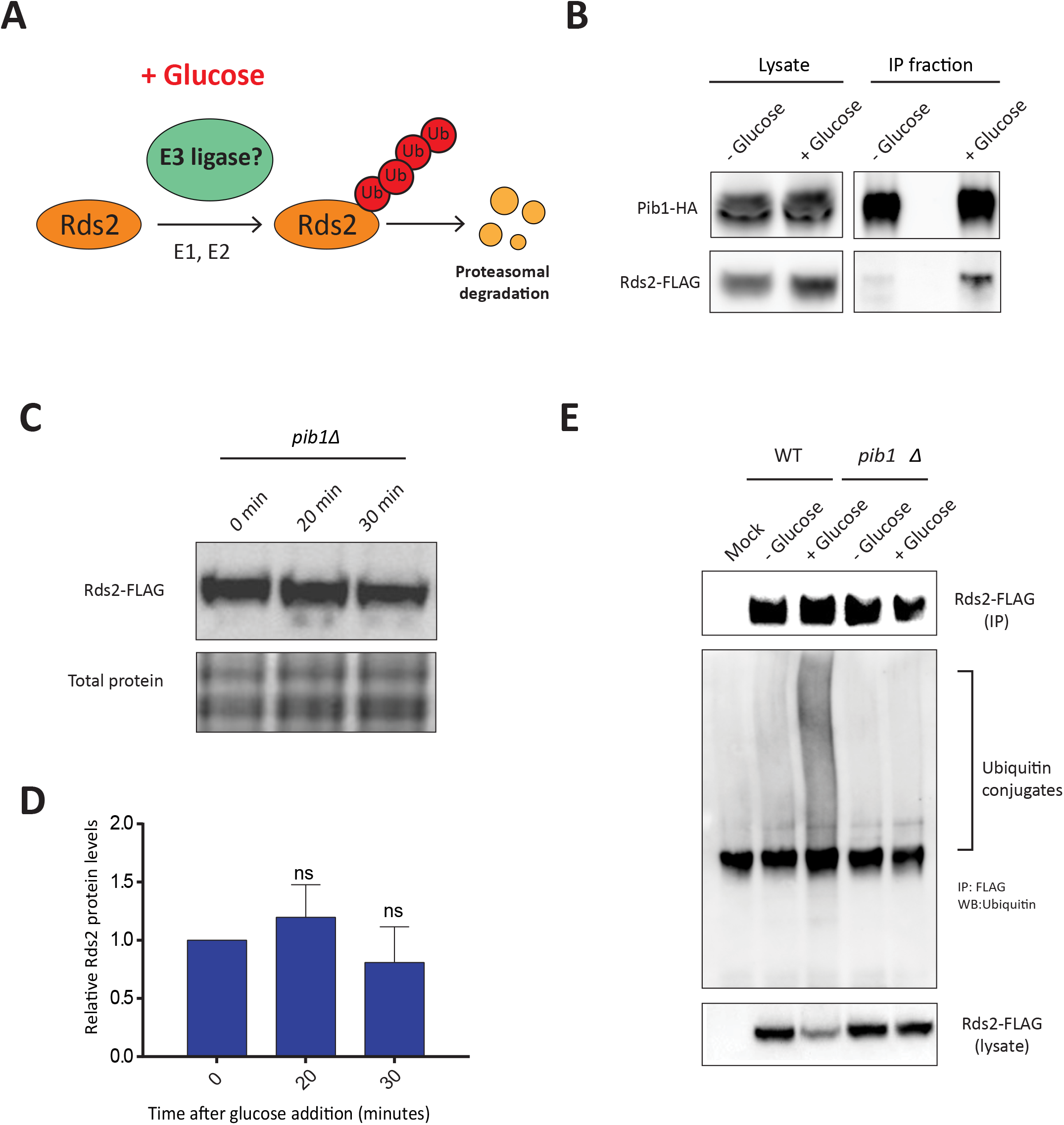
E3 ubiquitin ligase Pib1 interacts with and ubiquitinates Rds2. A) A schematic, illustrating a possible mechanism for glucose-mediated ubiquitination of Rds2, through an as yet unknown E3 ubiquitin ligase. B) The E3 ubiquitin ligase Pib1 interacts with Rds2 in a glucose-dependent manner. Pib1-HA was immunoprecipitated from glucose-treated and non-treated cells using an anti-HA antibody. The immunoprecipitated fractions were tested for the presence of Rds2-FLAG, using an anti-FLAG antibody, by Western blotting. C) Pib1 regulates glucose-mediated Rds2 degradation. Rds2-FLAG:*pib1Δ* cells grown in glucose-limited medium were treated with glucose, and Rds2 protein was measured at different time points by Western blotting. D) Quantification of the blot in (C) using ImageJ. Data are displayed as means ± SD, n=3. *p<0.05, **p0.01, ***p<0.001. E) Pib1 regulates glucose-mediated Rds2 ubiquitination. Rds2-FLAG was immunoprecipitated from glucose-treated and non-treated cells, separated on a 4-12% bis-tris gel, and Rds2-ubiquitin conjugates were detected by western blotting using an anti-ubiquitin antibody.

These data, therefore, suggested a possible involvement of Pib1 in regulating glucose-mediated Rds2 ubiquitination. To directly address this possibility, we first monitored Rds2 amounts after glucose addition, in cells lacking Pib1 (*pib1Δ*). Notably, Rds2 amounts remained stable and did not decrease, even ~30 minutes of glucose addition (Figures 3C and 3D). This was similar to the effect observed in cells treated with the proteasomal inhibitor MG132. Next, in order to determine whether Pib1 directly regulates Rds2 ubiquitination, we monitored the accumulation of poly-ubiquitin in Rds2 immunoprecipitates, in *pib1Δ* cells. Strikingly, no Rds2-polyubiquitin conjugates were observed even in the presence of glucose in *pib1Δ* cells (Figure 3E), in contrast to earlier observations made in wild-type cells. These data collectively show that Pib1 is required for the glucose-dependent ubiquitination and subsequent proteasomal degradation of Rds2 (Figure 3E).

### Pib1 ubiquitinates Rds2 *in vitro*

In order to determine if Pib1 can directly act as an E3 ligase that ubiquitinates Rds2, we reconstituted an *in vitro* ubiquitination assay system. This assay requires incubating the substrate protein (Rds2), ATP and ubiquitin with purified E1, E2 and E3 enzymes in a suitable reaction buffer, followed by western blotting with anti-ubiquitin antibody to detect ubiquitin conjugates of the substrate (Figure 4A). *S. cerevisiae* has a single E1 ubiquitin-activating enzyme-Uba1 (49). We also utilized the E2 ubiquitin-conjugating enzyme Ubc4, since it has been used as an E2 ligase with Pib1 in a previous study (50). We first expressed recombinant Uba1 and Ubc4 (with C-terminal GST tags) in *E.coli,* and subsequently purified these proteins (Figure 4B). These proteins were then utilized in predetermined amounts in the *in vitro* assay. Separately, we immunoprecipitated Rds2 (substrate) from *pib1Δ* cells, and Pib1 (E3-ligase) from *rds2Δ* cells treated with glucose, to obtain sufficient amounts of the substrate and the E3 ligase. BSA was used as a control protein in the reaction to test the substrate specificity of Pib1. Using this assay, with permutations of individual combinations of reagents (as indicated in Figure 4C), robust Rds2-ubiquitin conjugates were observed exclusively when the E1, E2, Pib1 and ubiquitin (along with ATP) were present in the reaction mixture (Figure 4C). Pib1 did not ubiquitinate the control protein (BSA) in these conditions, and the E1 and E2 enzymes alone could not ubiquitinate Rds2 (Figure 4C). These data show that Pib1 specifically, directly ubiquitinates Rds2 *in vitro.* Finally, in order to understand the glucose specificity of this process, we performed this *in vitro* ubiquitination assay using Rds2 (substrate) immunopurified from cells growing in glucose-limited conditions, with or without glucose addition for ~20 minutes. Notably, no Rds2-ubiquitin conjugates were observed with Rds2 immunoprecipitated from cells withtout glucose addition, while clear poly-ubiquitination was observed in Rds2 immunopurified from cells post glucose-addition (Figure 4D). This confirms that Pib1 mediated Rds2 ubiquitination is glucose dependent. Collectively, these data establish Pib1 as the E3 ubiquitin ligase that ubiquitinates Rds2 upon glucose re-availability, leading to Rds2 degradation.

**Figure 4:**
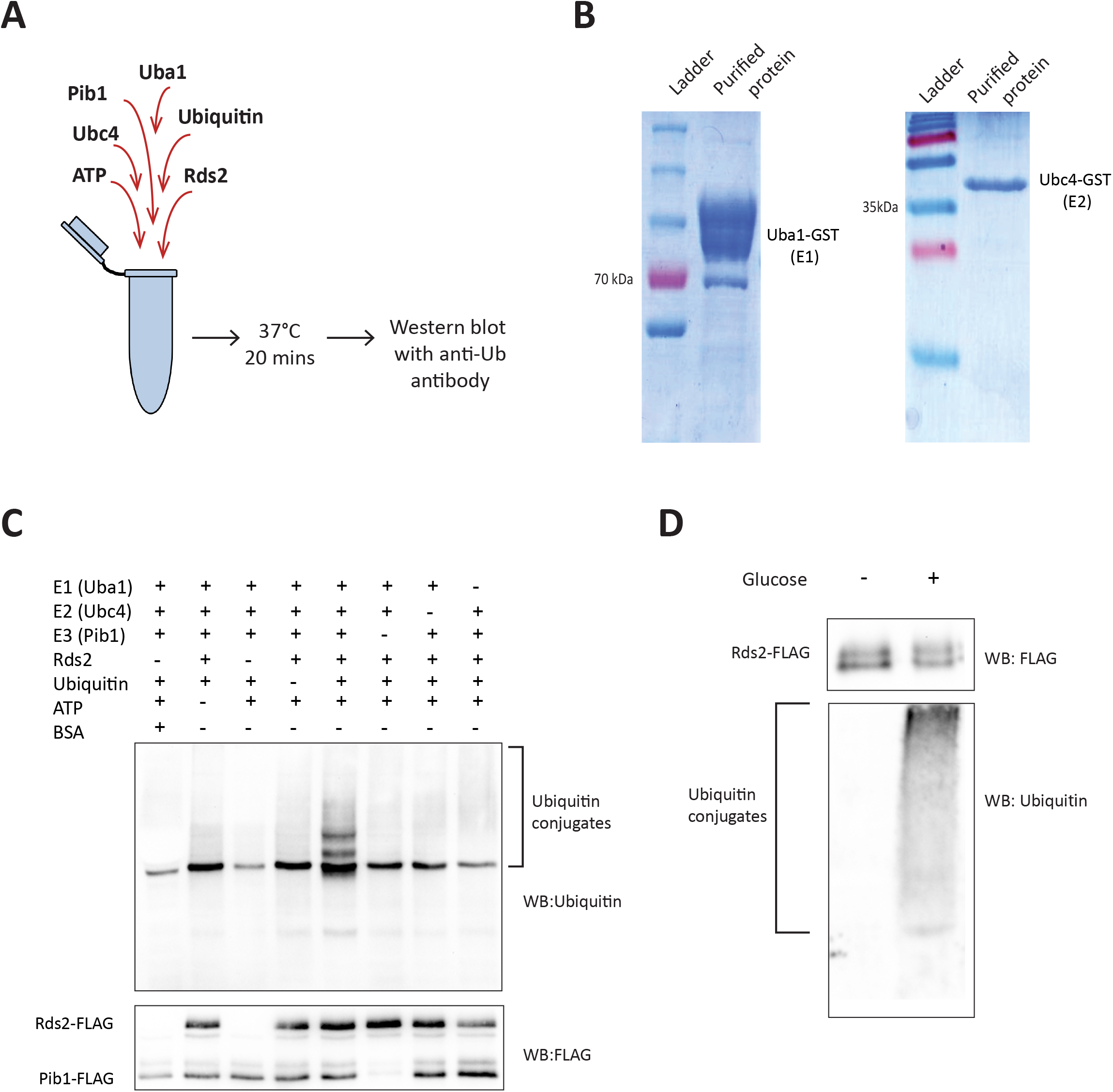
Pib1 ubiquitinates Rds2 *in vitro*. A) A schematic illustration of the *in vitro* ubiquitination assay design, to test for the ability of Pib1 to specifically ubiquitinate Rds2 *in vitro.* B) Purification of recombinant E1 and E2 enzymes. GST tagged E1 (Uba1) and E2 (Ubc4) enzymes were recombinantly expressed in BL21 (DE3) cells, and purified by glutathione affinity chromatography. The purified fractions run on SDS-PAGE are shown. C) Pib1p ubiquitinates Rds2p *in vitro.* An *in vitro* ubiquitination assay was set up with the following components in the reaction mixture-E1 (Uba1p,) E2 (Ubc4p), E3 (Pib1p-Flag), the substrate (Rds2p-Flag, or bovine serum albumin-BSA as a control), ubiquitin, MgCl_2_ and ATP. The mixture was incubated for 20 minutes at 37°C and the reaction was terminated by adding SDS-PAGE sample buffer. The mixture was run on a 10% SDS-PAGE gel and western blotting was done using anti-FLAG (to detect Rds2-FLAG and Pib1-FLAG), or anti-Ubiquitin (to detect Rds2-ubiquitin conjugates) antibodies. D) Pib1 mediated Rds2 ubiquitination is specific to the presence of glucose. An *in vitro* ubiquitination assay was performed using Rds2 immunoprecipitated from glucose-limited cells, either before glucose addition or 20 minutes after glucose addition, to determine if ubiquitination of Rds2 by Pib1 was glucose dependent. The samples were run on a 10% gel and ubiquitin conjugates were detected by western blotting with anti-ubiquitin antibody.

### Pib1 mediates the effective shutdown of gluconeogenesis following glucose addition

If Pib1 is important for overall adaptation of cells to the presence of glucose (i.e. glucose repression), cells lacking Pib1 could be expected to show some defects in fully shutting down gluconeogenesis, after glucose re-entry. Therefore, since Rds2 transcriptionally activates the gluconeogenic enzyme genes – *FBP1* and *PCK1,* cells lacking Pib1 may accumulate these transcripts after glucose addition. To test this, we first estimated transcript levels of these genes in WT and *pib1Δ* cells at different time points after glucose addition by qRT-PCR. We observed significantly higher mRNA levels of both *FBP1* and *PCK1* in *pib1Δ* cells ~one hour post glucose addition, compared to the WT cells (Figure 5A). Finally, we tested the functional implications of the loss of Pib1, and inefficient shutting down of the gluconeogenic machinery (after glucose addition), using a competitive growth fitness assay with WT and *pib1Δ* cells. WT and *pib1Δ* cells (carrying different drug selection markers) were grown separately in glucose-replete, or glucose-limited medium, and equal numbers of these cells (based on absorbance at OD_600_) were mixed together in either glucose-replete or glucose-limited medium. Cells were plated on drug selection plates at different time intervals, and the relative number of WT and *pib1Δ* were calculated by colony counting. This experimental design is illustrated in Figure 4B. Under conditions when cells are transferred from glucose poor to glucose replete medium, the *pib1Δ* cells will not as efficiently switch to glycolysis (i.e. show efficient glucose repression) as WT cells, predicting that WT cells will outcompete *pib1Δ* cells in growth. Indeed, we observed that the relative number of WT cells is higher compared to *pib1Δ* cells after the switch to a glucose-replete medium, for around 4.5 hours (Figure 4B). Eventually, over longer times, the *pib1Δ* cells catch up with the WT cells. Note: in control experiments (where the cells were grown and mixed in glucose-replete medium), both WT and *pib1Δ* cells showed equal numbers of cells (Figure 4B). Also interestingly, when cells were grown and mixed together in glucose-deficient (glycerol/ethanol) medium, *pib1Δ* cells outcompeted WT cells around 6 hours after cells were shifted to this medium (Figure 4B), further supporting a role for Pib1 in regulating a gluconeogenic state. Collectively, these data suggest that Pib1 regulates the ability of cells to efficiently switch to a glucose-repressed state, by controlling the amounts of the gluconeogenic regulator Rds2, upon glucose re-entry.

**Figure 5:**
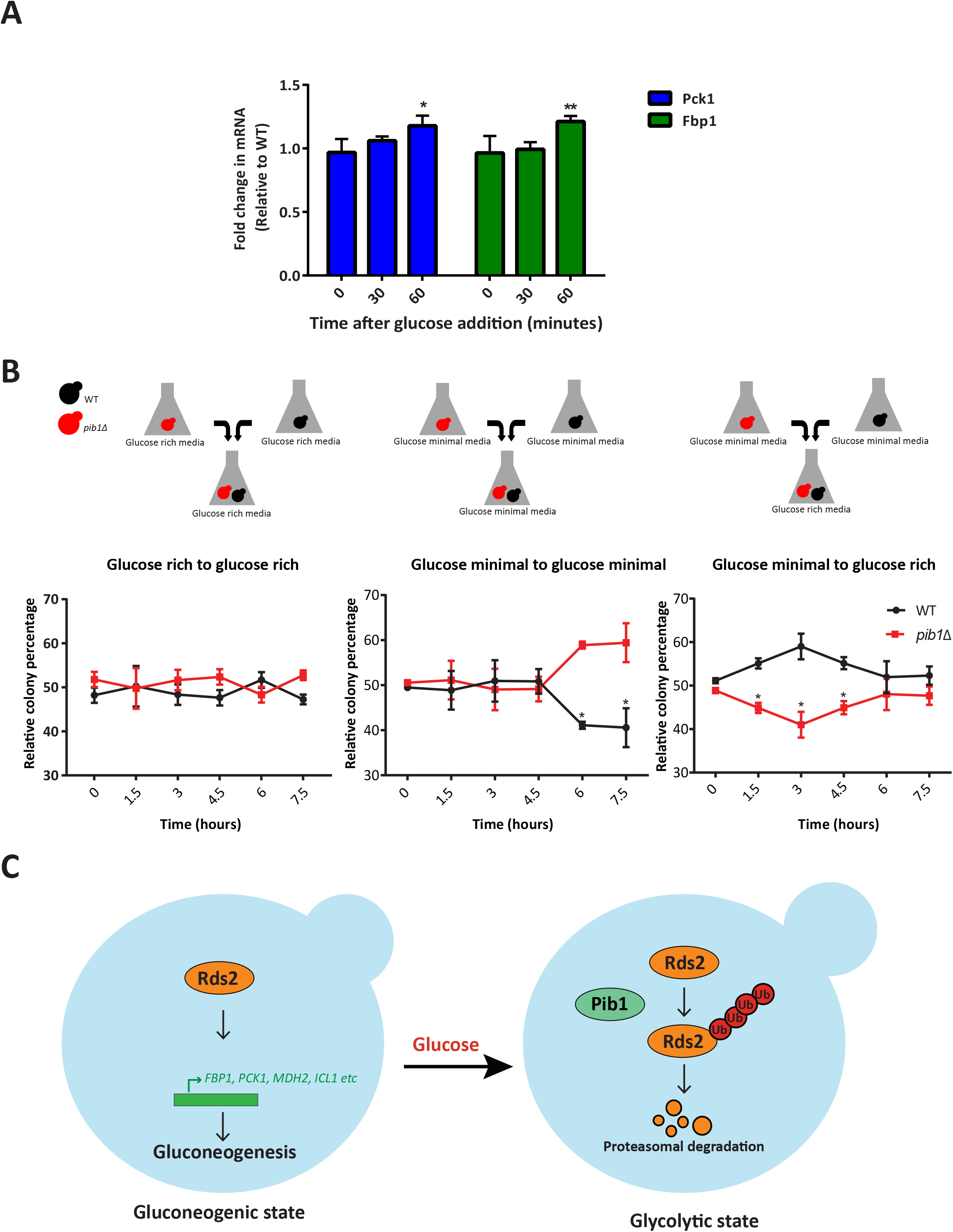
Pib1 regulates the effective shutdown of gluconeogenesis following glucose addition. A) *PCK1* and *FBP1* mRNAs are higher in *pib1Δ* cells, compared to WT cells, after glucose addition. The mRNA levels of *PCK1* and *FBP1* were measured using qRT PCR in WT and *pib1Δ* cells, at different time points after glucose addition. Fold changes in mRNA amounts are presented. Data are displayed as means ± SD, n=3. *p<0.05, **p0.01, ***p<0.001. B) *pib1Δ* cells show a fitness defect in competitive growth assays upon glucose addition. WT and *pib1Δ* cells were cultured in either glucose-replete or glucose-limited medium, and equal amounts of cells were mixed together, and transferred to glucose-replete medium, glucose-limited medium (2% glycerol/ethanol), or glucose-limited medium supplemented with glucose (final concentration of 3%), with a starting cell density based on OD_600_ of 0.2 each. This experimental design is illustrated. Subsequently, cells were collected and plated on appropriate selection plates, to quantify relative amounts of each genetic background. The relative genetic background is shown. Data are displayed as means ± SD, n=3. *p<0.05, **p0.01, ***p<0.001, significance is based on a student’s T-test. C) A proposed model illustrating the role of Pib1 in regulating glucose-mediated Rds2 degradation. In glucose-limited cells, Rds2 transcriptionally activates the expression of different gluconeogenic and glyoxylate cycle enzymes, enforcing a gluconeogenic state. Upon glucose addition, Pib1 interacts with and ubiquitinates Rds2, targeting it for proteasomal degradation.

## Discussion

In this study, we identify a functional role for the E3 ubiquitin ligase Pib1, as a mediator of effective gluconeogenic shutdown upon glucose re-entry into the medium. Pib1 binds to and mediates the glucose-dependent ubiquitination, and subsequent proteasomal degradation of the gluconeogenic transcription factor Rds2. Competitive growth experiments suggest a role for Pib1 in mediating effective glucose repression, suggesting a role for Pib1 in mediating an ‘off-switch’ for gluconeogenesis.

We first characterized glucose-mediated Rds2 regulation, and using this as our readout, our aim was to identify regulators involved in downregulating the gluconeogenic transcription factors, when glucose becomes available to cells. Upon glucose entry, Rds2 protein amounts rapidly decrease over time (Figure 1). This glucose-mediated decrease in Rds2 amounts is due to its ubiquitin-mediated proteasomal degradation (Figure 2). While other studies have identified regulators gluconeogenic and glyoxylate cycle enzymes themselves, our study identifies a regulator of the transcription factor that controls gluconeogenesis shut-down upong glucose re-entry, and therefore the overall transcriptional state of the cell with changing nutrients.

Through this, we identify a glucose-dependent, ubiquitin-mediated signaling response, and a role for the E3 ubiquitin ligase Pib1 in this response. Strikingly, the interaction of Pib1 with Rds2 is very specific to the re-availability of glucose (Figure 3). Further, the loss of *PIB1* prevents Rds2 ubiquitination and its proteasomal degradation even in the presence of glucose (Figure 3). This interaction of Pib1 with Rds2, and its role in ubiquitinating Rds2 is direct, and glucose-dependent (Figure 4). Finally, we find that functionally, Pib1 regulates both the amounts of gluconeogenic enzyme transcripts (targets of Rds2), and cell growth and competitive fitness post glucose addition (Figure 5).

Collectively, these observations reveal a novel, functional role for Pib1 as a regulator of the glucose-mediated ubiquitination and proteasomal degradation of Rds2, and therefore effective glucose repression in yeast. While previous studies have suggested the role of Pib1 in regulating vacuolar sorting (51), and a recent study has identified the exocyst subunit Sec3p as a Pib1 specific target in *S. pombe* (52), no function of Pib1 related to regulating responses to nutrients is currently known.

These results add a dimension to recent studies that suggest roles of E3 ubiquitin ligases as regulators of metabolic states (Nakatsukasa et al., 2015; Santt et al., 2008). The glucose-dependent Pib1 mediated signaling event which we identified takes place within minutes after glucose addition, reiterating the swiftness of the response. Since E3 ligases can ubiquitinate specific sets of substrates, an exciting area of investigation will be to identify substrates of Pib1, in a glucose-dependent context.

The ubiquitination machinery is itself regulated at multiple levels. Apart from E3 ligases which conjugate ubiquitin to specific targets, the deubiquitination machinery removes ubiquitins from target proteins (54, 55). This involves deubiquitinase enzymes that bind to specific ubiquitinated proteins and cleaves the bound ubiquitins, to co-ordinately achieve a tight, reversible regulation of protein ubiquitination. The role of deubiquitinase enzymes in metabolic state switching has largely remained entirely unexplored. Our experimental approach, to observe early ubiquitination responses to the re-entry of nutrients, might reveal additional such regulators.

Finally, a less-explored, but an interesting signaling possibility is the cross-talk of ubiquitin machinery with other post-translational modifications, notably acetylation and phosphorylation (56, 57). How this cross-talk between PTMs activates or inhibits target protein ubiquitination, in the context of different nutrient cues, is poorly explored. Emerging studies suggest that acetylation and phosphorylation can regulate the ubiquitination and degradation of metabolic enzymes in other model systems (58). The combinatorial effect of these PTMs could, therefore, result in a highly sensitive, tightly controlled, rapidly responsive metabolic switching regulation machinery. Thus, the complex network of metabolic state regulation machinery involving E3 ubiquitin ligases, deubiquitinases, kinases, phosphatases, acetylases, and deacetylases opens an interesting arena of post-translational modification mediated regulation of metabolic switching.

## Experimental procedures

### Yeast strains, media and growth conditions

The prototrophic CEN.PK strain (WT) was used in all experiments (59). All the strains used in this study are listed in Table S1. The gene deletion and C-terminal chromosomal tagging was done using the standard PCR based gene deletion and modification strategy (60). The media used in this study were glucose-limited media (1% Yeast extract, 2% peptone, 2% ethanol 10 and 1% glycerol) and glucose-replete media (1% Yeast extract, 2%peptone, and 2% dextrose). For experiments involving the addition of glucose to cells grown in glucose-limited media, cells were initially grown at 30°C to an OD600 of 0.8 and glucose was added to a final concentration of 3%. All experiments involving MG132 were done in a *pdr5Δ* background to prevent the efflux of MG132 by the multidrug transporter protein Pdr5. A final concentration of 100 μM MG132 was used in all the experiments involving proteasomal inhibition.

### Protein extraction and western blotting

When the cells reach an OD600 of 0.8, 10 ml of the culture cells were collected, pelleted and protein extraction was performed using trichloroacetic acid (TCA) extraction method. Briefly, the pelleted cells were resuspended in 400 μl 10 % TCA solution and the cells were lysed by bead beating (3 times, 20 seconds each with 1 min cooling in ice-water slurry between each cycle). Lysate extracts were centrifuged and the protein pellets thus obtained were resuspended in 400 μl SDS-glycerol buffer (7.3% SDS, 29.1% glycerol and 83.3 mM Tris base), and heated for 10 minutes at 100°C. The supernatant was collected after centrifugation (20000g/10 min), and total protein estimation was done using BCA assay (BCA assay kit, G-Biosciences). Samples were normalized (to maintain constant protein amounts) in SDS-glycerol buffer. The samples were run on 4-12 % Bis-Tris gels (Invitrogen) unless specified. Western blots were developed using the following antibodies – anti-FLAG M2 (mouse mAb, Sigma), anti-HA (12CA5 mouse mAb, Roche), anti-ubiquitin (P4D1 mouse mAb, CST) and anti-FLAG (D6W5B rabbit mAb, CST). Horseradish peroxidase-conjugated secondary antibodies (mouse and rabbit) were obtained from Sigma. For developing the blot, standard enhanced chemiluminescence reagent (WesternBright ECL HRP substrate, Advansta) was used. Coomassie-stained gels were used as the loading controls. The quantification of the blots was done using ImageJ. The statistical significance was determined using Student’s t-test (using GraphPad Prism 7.0)

### Immunoprecipitation and co-immunoprecipitation

Cells were grown in glucose-limited media with or without glucose addition as specified and 50 OD_600_ of cells were harvested by centrifugation. The collected cells were lysed in lysis buffer (50 mM HEPES buffer pH-7.5, 50 mM Sodium fluoride, 10% Glycerol, 150 mM KCl, 1 mM EDTA, 2 mM Sodium orthovanadate, 2 mM PMSF, 0.1 mM Leupeptin, 0.02 mM Pepstatin, 0.25% Tween 20) using a mini bead beater. The lysate was incubated with Dynabeads Protein G (Invitrogen), conjugated with the required antibody, for 3 hrs at 4°C. The lysate was collected and the beads were washed 5 times in wash buffer (50 mM HEPES buffer pH-7.5, 50 mM Sodium fluoride, 10% Glycerol, 150 mM KCl, 1 mM EDTA, 2 mM Sodium orthovanadate, 0.05% Tween 20). The bound proteins were eluted by boiling the samples in SDS-PAGE sample buffer (for HA-tagged proteins), or by FLAG peptide elution (for FLAG-tagged proteins, using 0.5 mg/ml FLAG peptide). The samples were then subjected to western blotting. For co-immunoprecipitation studies with Pib1 and Rds2, HA-tagged Pib1 was immunoprecipitated using the above-described method using and eluted by boiling in SDS-PAGE sample buffer. Rds2-FLAG was detected in the elution fraction by western blotting using anti-FLAG antibody.

### *In vivo* ubiquitination assay

Rds2-FLAG was immunoprecipitated from glucose-treated and non-treated cultures using anti-FLAG M2 antibody (mouse mAb, Sigma) and eluted by FLAG peptide elution. The eluted proteins were run on a 10% polyacrylamide gel and subjected to western blotting. The presence of Rds2-FLAG in the lysate and elution fractions was detected using anti-FLAG antibody (D6W5B rabbit mAb, CST). In order to detect Rds2-ubiquitin conjugates, anti-ubiquitin antibody (P4D1 mouse mAb, CST) was used.

### *In viro* ubiquitination assay

Uba1-GST and Ubc4-GST were expressed in E.coli BL21(DE3) cells using pGST parallel plasmids, and purified using glutathione-affinity chromatography. Briefly, the lysate was passed through a column containing glutathione resin. The bound proteins were eluted using an elution buffer containing 10 mM reduced glutathione. The GST tags were removed using 6X His tagged TEV protease treatment, followed by passing it through a column containing glutathione resin. The TEV protease was removed by passing this sample through a Ni-NTA column. The concentrations of the purified proteins were estimated using Bradford estimation method. Rds2-FLAG was immunoprecipitated from *pib1Δ* cells and Pib1-FLAG from *rds2Δ* cells and eluted by FLAG peptide elution. The concentrations of these proteins were estimated by BCA protein estimation method (BCA assay kit, G-Biosciences). The *in vitro* ubiquitination assay was set up with the following components in the reaction mixture-5 nM Uba1p, 100 nM Ubc4p, 20 nM Pib1-FLAG, 200 nM Rds2-FLAG, 0.02 mg/ml ubiquitin, 1X ubiquitination buffer (50 mM Tris-HCl pH-8, 5 mM MgCl2, 0.1% Tween 20, 1 mM DTT), 2 mM ATP. The mixture was incubated for 20 minutes at 37°C and the reaction was terminated by adding SDS-PAGE sample buffer. The samples were run on a 10% polyacrylamide gel and ubiquitin chains were detected by western blotting with anti-ubiquitin antibody (P4D1 mouse mAb, CST).

### RNA isolation and RT-qPCR

RNA isolation was done using a hot acid phenol extraction method. Briefly, cells were grown in glucose-limited media and 5 OD_600_ cells were harvested by centrifugation. The pellets were resuspended in 400 μl TES solution (10 mM Tris-HCl pH 7.5, 10 mM EDTA and 0.5% SDS). To this, 400 μl acid phenol was added and mixed by vortexing. The resuspended pellets were incubated at 65°C for 60 minutes with intermittent vortexing. The supernatant was collected after centrifugation and subjected to another round of acid phenol treatment followed by incubation and intermittent vortexing as before. The supernatant obtained by centrifugation was treated with 400 μl chloroform, mixed and the aqueous phase obtained after centrifugation was collected. To this, 40 μl 3M sodium acetate, pH 5.3 and 1 ml ice-cold 100 % ethanol was added. This mixture was incubated in −20°C for 60 minutes. RNA was pelleted by centrifugation and washed in ice-cold 70 % ethanol. The pellet was air dried and resuspended in nuclease-free water. The isolated RNA was quantified and treated with Turbo DNase (Invitrogen). DNase treated RNA was then used for cDNA synthesis using Superscript III reverse transcriptase (Invitrogen) as per the manufacturer’s protocol. The cDNA was then used to perform qRT PCR using the KAPA SYBR FAST qRT PCR master mix kit (KK4602, KAPA Biosystems) as per the manufacturer’s instructions. Actin *(ACT1)* was used as a control to normalize the values obtained. The mRNA fold change was calculated by a standard 2^-(ddct) method. The statistical significance was determined using Student’s t-test (using GraphPad Prism 7.0).

### Competitive growth fitness assay

WT and Pib1Δ cells carrying different drug selection markers were cultured separately in either glucose-replete or glucose-limited medium. At an OD600 ~ 0.8, they were subcultured together at a starting OD600 of 0.2 each, in either glucose-replete medium, glucose-limited medium or glucose-limited medium supplemented with glucose (final concentration of 3%). Samples were collected from this mixed culture at 0 hr, 1.5 hrs, 3 hrs, 4.5 hrs, 6 hrs, and 7.5 hrs and were serially diluted to a final concentration of ~ 3000 cells per ml. 100 μ1 of this was plated on appropriate drug selection plates in triplicates and incubated at 30°C. The number of colonies was counted and the relative percentage of WT and Pib1Δ colonies at each time point was calculated. These values obtained were plotted and the statistical significance was determined using Student’s t-test (using GraphPad Prism 7.0).

## Acknowledgements

VV is supported by a DST-INSPIRE fellowship (DST/INSPIRE/03/2016/001546) from the Department of Science and Technology, Govt. of India. SL acknowledges support from a Wellcome Trust-DBT India Alliance Intermediate Fellowship (IA/I/14/2/501523), and institutional support from inStem and the Dept. of Biotechnology, Govt. of India.

## Supplementary Figure legends

**Supplementary Figure 1:**
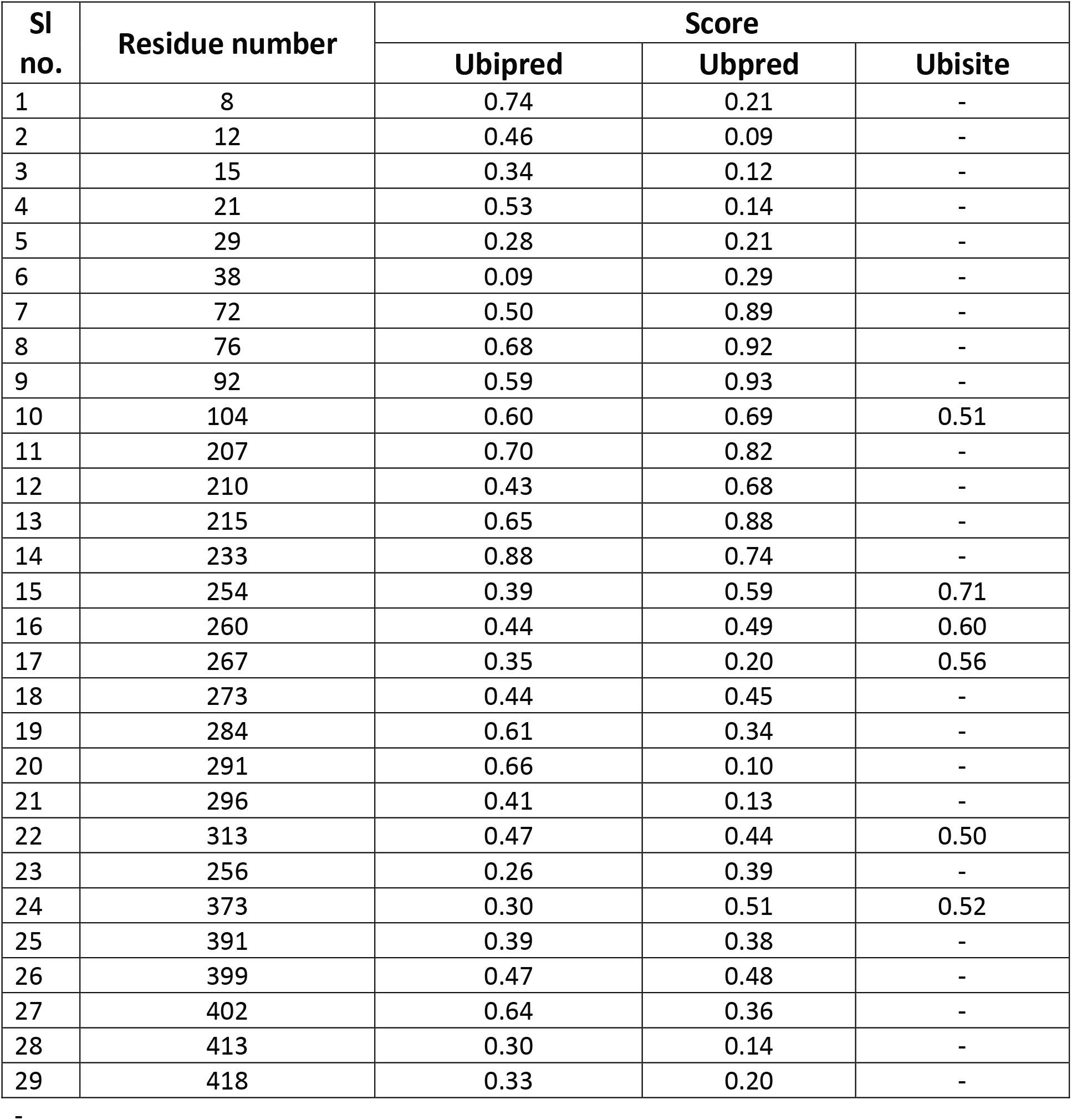
Lysine residues in Rds2, with their predicted scores for ubiquitination. Multiple prediction programs (Ubpred, Ubisite, and Ubipred) were used to find out the probable ubiquitination sites in Rds2. The lysine residues present in Rds2, along with the scores for ubiquitination, obtained from the predictions are indicated.

**Supplementary Figure 2:**
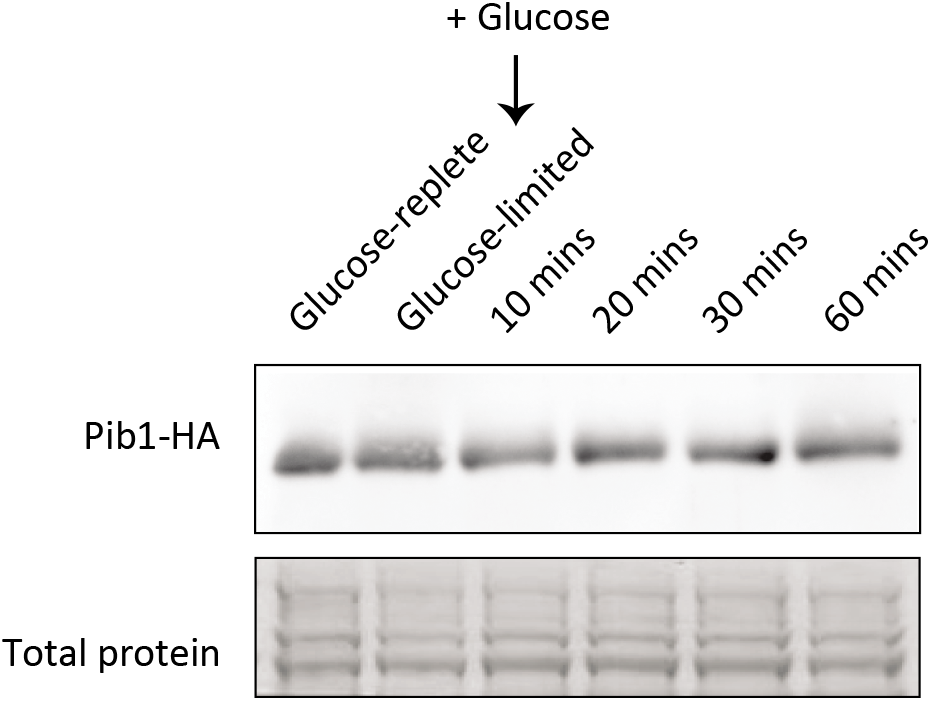
Pib1 protein before and after glucose addition to cells in glucose-limited medium. Pib1 protein amounts were analyzed by Western blotting, using samples collected from glucose-limited medium, before and at different time points after glucose addition. No change in the protein levels was observed. Coomassie-stained gels were used as loading control.

